## Supplementary material for "Higher-order connectomics of human brain function reveals local topological signatures of task decoding, individual identification, and behavior": SI

Andrea Santoro[1,2], Federico Battiston[3], Maxime Lucas[4], Giovanni Petri[4,5,6], Enrico Amico[1,2]
[1]Neuro-X Institute, EPFL, Geneva, Switzerland
[2]Department of Radiology and Medical Informatics, University of Geneva, Geneva, Switzerland
[3]Department of Network and Data Science, Central European University, 1100 Vienna, Austria
[4]CENTAI Institute, 10138 Torino, Italy
[5]NPLab, Network Science Institute, Northeastern University London, London, E1W 1LP, United Kingdom
[6]Networks Unit, IMT Lucca Institute, 55100 Lucca, Italy

December 4, 2023

### S1 Task decoding

In the main text, we have shown the task decoding ability of the four methods on resting-state and task fMRI data of the HCP dataset. In particular, once the time-time correlation matrices have been computed, we binarize the matrices by considering a common threshold at the $95th$ percentile. While this choice provides the highest task decoding ability in terms of the element-centric similarity (ECS) by all methods, we report in Fig. S1-S2 similar plots when considering two other thresholds, namely, $90th$ and $97th$ percentiles.

### S2 Brain Fingerprinting

As a second application, we have also considered functional brain fingerprinting [2, 1]. That is, the ability to identify a subject from a group, solely based on their brain's distinctive functional pattern.

We compared the four different methods on the HCP dataset relying on two sessions of resting-state fMRI data (i.e., sessions $REST\_1\_LR$ and $REST\_2\_LR$ for test and retest, respectively) by assessing the quality of fingerprinting using the differential identifiability measure [1]. We report in Fig. S3 the performances of the functional brain fingerprinting across the four methods considered to analyze resting-state fMRI data. In particular, when focusing on whole-brain connections we find similar performances across methods. The performance scores are in line with some previous results [1], but can slightly differ across studies [3] given that identifiability scores are typically influenced by different steps of the fMRI preprocessing, or brain parcellation considered (in this study we considered 19 ROIS among subcortical and cerebellum, for a total of 119 ROIs).

As done in the main text, we can also explore subject-specific patterns of brain activation. We report in Fig. S4 the coefficient of variation (cv) for the triangle approach, when projecting the triangle weights of connections that involve either one (respectively two and three) nodes within one of the seven resting-state functional networks on the cortical brain surface, averaged across the 100 HCP subjects. Interestingly, the interactions between unimodal (i.e. visual, somatosensory) areas and transmodal (e.g. DMN, frontoparietal) brain areas significantly vary between subjects, as indicated by the high values of the coefficient of variation between the seven functional networks.

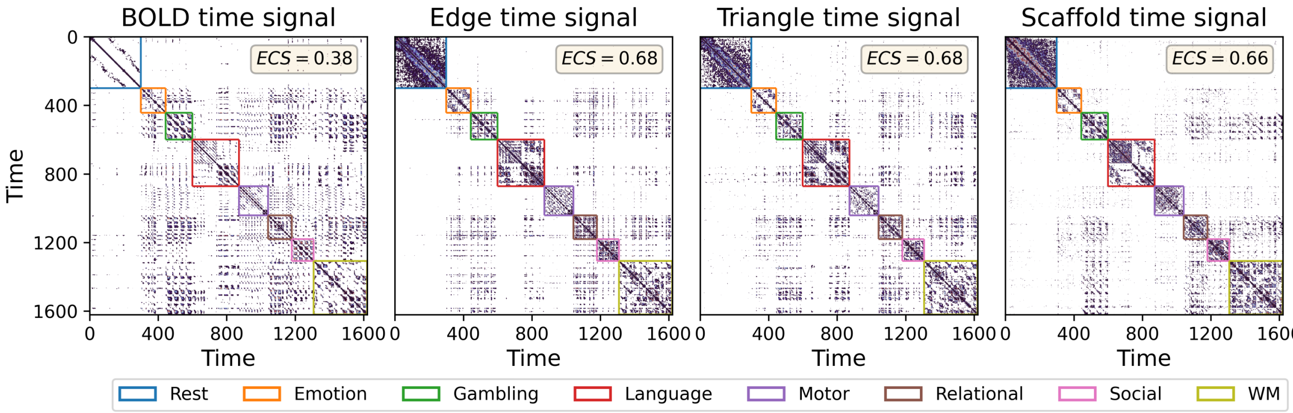


Figure S1: **Local higher-order topological indicators for fMRI task differentiability at** $90th$ **percentile**. As done in the main text, we compute *local* observables by comparing the temporal recurrence plots (i.e. time-time correlation matrices) for the four methods, from the lower-order methods (BOLD and edge signals) to higher-order ones (triangle and scaffold time signals). We set a common threshold at the $90th$ percentile to binarize the data when analyzing an fMRI temporal signal obtained by concatenating resting-state and seven HCP tasks. Colored boxes within the plots denote the ground truth of rest and task segments. The element-centric similarity (ECS) measure provides a quantification of the task decoding ability of the different methods. Here, edges and triangles lead the pack, yet with a lower value than the one reported in the main text. Results are averaged over the 100 subjects, considering the mean between LR and RL phase encoding.

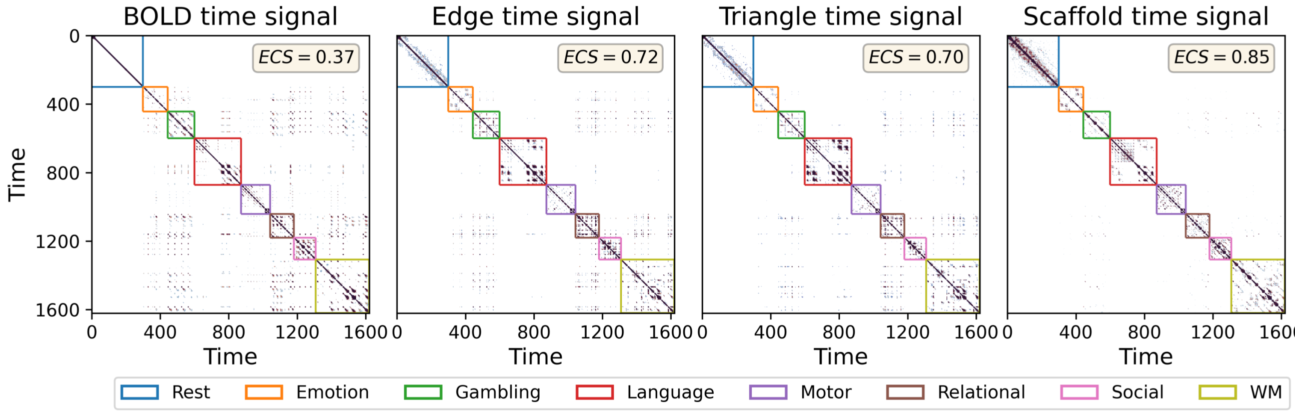


Figure S2: **Local higher-order topological indicators for fMRI task differentiability at** $97th$ **percentile**. As done in the main text, we compute *local* observables by comparing the temporal recurrence plots (i.e. time-time correlation matrices) for the four methods, from the lower-order methods (BOLD and edge signals) to higher-order ones (triangle and scaffold time signals). We set a common threshold at the $97th$ percentile to binarize the data when analyzing an fMRI temporal signal obtained by concatenating resting-state and seven HCP tasks. Colored boxes within the plots denote the ground truth of rest and task segments. The element-centric similarity (ECS) measure provides a quantification of the task decoding ability of the different methods. Here, the scaffold method leads the pack, yet with a lower value than the one reported in the main text. Results are averaged over the 100 subjects, considering the mean between LR and RL phase encoding.

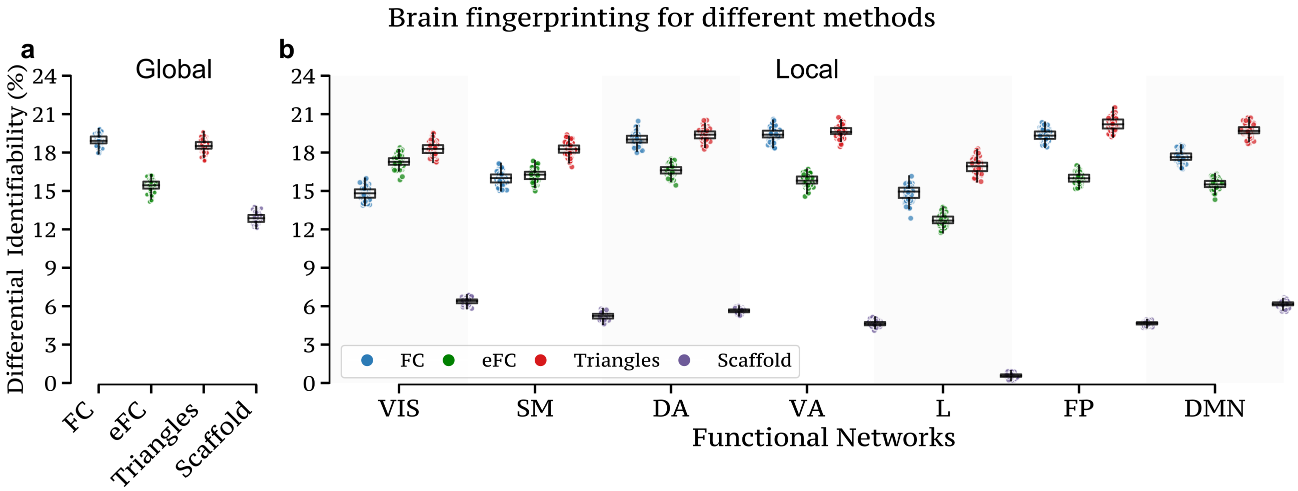


Figure S3: **Global and local functional brain fingerprinting performance across methods**. (**a**) We report the differential identifiability scores obtained when focusing on whole-brain connections. In this scenario, the identifiability scores of the four methods exhibit no significant differences, with all methods yielding very similar results, with FC and triangles leading the pack. (**b**) When repeating the analysis considering only specific local connections, we also report the scores obtained when considering the functional connections having at least one node in the functional network analyzed, namely, visual (VIS), somatomotor (SM), dorsal attention (DA), ventral attention (VA), limbic (L), frontoparietal (FP), and default mode network (DMN).

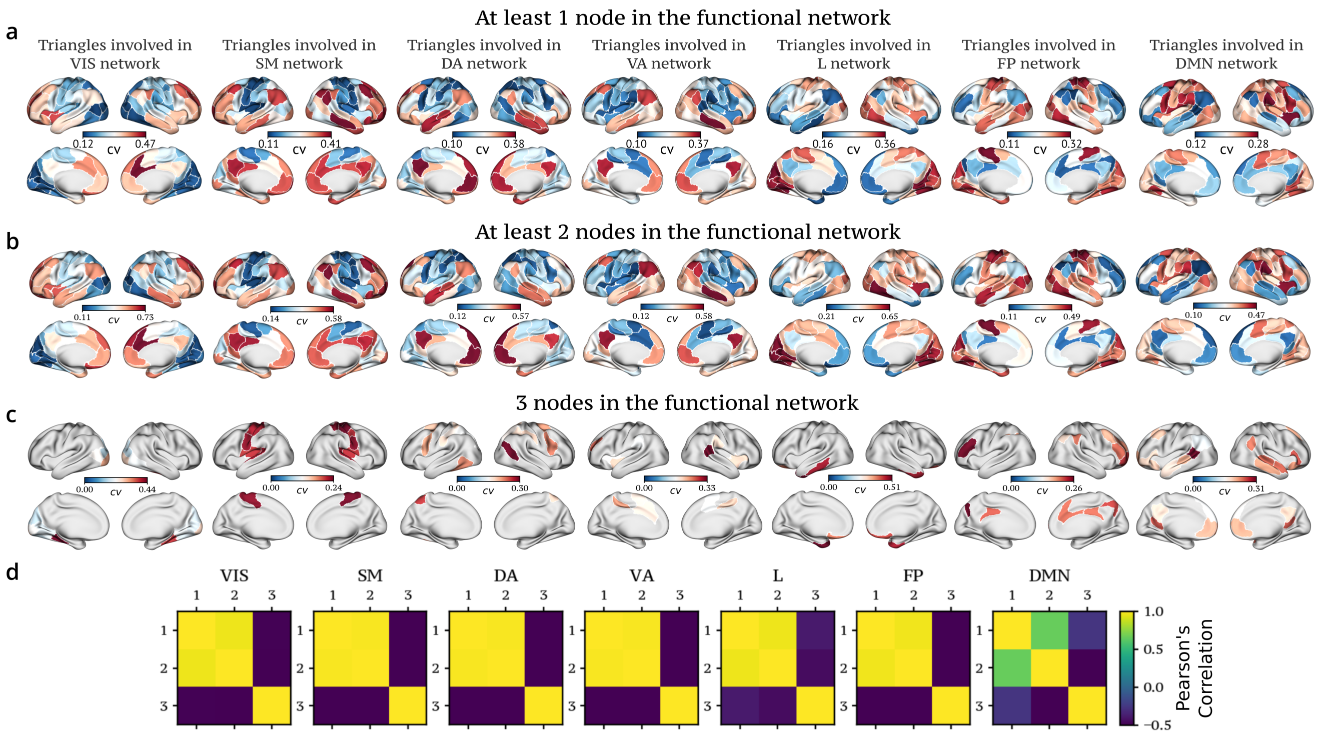


Figure S4: **Cortical brain projections for functional brain fingerprinting**. **(a)** We report the subject-specific brain activation patterns by examining the coefficient of variation (cv) for triangle nodal strength on the cortical brain surface, averaged across the 100 HCP subjects. Specifically, our focus extends to triangles having at least one **(a)**, two **(b)**, or three **(c)** nodes within the functional network of interest. As discussed in the main text, the interaction of somatosensory areas with higher-order networks (i.e., FP and DMN) exhibits considerable variability among subjects, as evidenced by the high values of the coefficient of variation. **(d)** The Pearson's correlation coefficient $\rho$ obtained from comparing cortical maps reveals distinctions only in the way triangle interactions occur within DMN and VIS networks.
